# The distortion-push mechanism for the γ-subunit rotation in F1-ATPase

**DOI:** 10.1101/2025.02.03.636200

**Authors:** Masahiro Motohashi, Mao Oide, Chigusa Kobayashi, Jaewoon Jung, Eiro Muneyuki, Yuji Sugita

## Abstract

F_1_-ATPase comprises the stator ring consisting of α₃β₃ subunits and the rotor γ subunit. The γ subunit rotation mechanism has been extensively investigated by biochemical analyses, structural studies, single-molecule measurements, and computational studies. Recent cryo-electron microscopy (cryo-EM) structures of F_1_-ATPase from the thermophilic bacterium *Bacillus* PS3 (TF_1_) provide us with further possibilities for a better understanding of the γ-rotation mechanisms. Using cryo-EM structures having the γ-rotation angles close to the binding dwell and catalytic dwell states, we investigate the relationships between the γ subunit rotation, conformational changes of the stator α_3_β_3_ subunits, and the nucleotide-binding and release. We performed targeted molecular dynamics (MD) simulations with external forces on the α₃β₃ subunits and observed 80° substep rotations of the γ subunit. Then, we optimized the most probable transition pathway through the mean-force string method simulations with 64 images. Finally, using umbrella sampling, we calculated the potential of mean forces along the minimum free energy pathway during the 80° substep rotation. Our MD simulations suggest that 80° substep rotation is divided into the first rotation, resting, and the second rotation. Notably, the first rotation is driven by the distortion of the stator α_3_β_3_ subunits, and the second rotation is induced mainly by direct β/γ subunit interactions. This model, which we call the distortion-push mechanism, is consistent with the residue-level experimental analysis on F_1_-ATPase and the atomic structures determined by X-ray crystallography and cryo-EM.

## Introduction

ATP, a primary energy source for many cellular activities, is synthesized by the ATP synthase, which is localized in the inner mitochondrial membrane, thylakoid membranes of chloroplasts, and the plasma membrane of bacteria. The ATP synthase consists of two rotary molecular motors: the proton-driven F_o_ and the ATP-driven F_1_. The F_o_ complex (ab_2_c_8-15_) functions through passive proton transports, resulting in the clockwise rotation of the c-ring when viewed from the cytoplasmic side. In contrast, the F_1_ complex (α_3_β_3_γδε) rotates counterclockwise during ATP hydrolysis (1). These two motors with opposite rotational directions interact through their respective stators and rotors, functioning cooperatively for their biological activities. When the clockwise rotation of F_o_ predominates, it transports protons accumulated in the electron transport chain to enable ATP synthesis by F_1_. This reaction is reversible; when the counterclockwise rotation of F_1_-ATPase dominates, F_1_ hydrolyzes ATP, and F_o_ expels protons against the concentration gradient (2, 3).

The isolated F_1_ is known to function as a rotary molecular motor in the form of the α₃β₃γ complex (F_1_-ATPase), which excludes the ε and δ subunits (1) (Fig. 1A). F_1_-ATPase comprises the stator ring consisting of α₃β₃ subunits and the rotor γ subunit. The rotor γ subunit rotates counterclockwise by hydrolyzing ATP at the three catalytic sites of the stator. The stator is a hexamer formed by assembling three heterodimers of the α and β subunits, each possessing an ATP catalytic site. Most catalytic residues are in the β subunits, which exhibit more flexible structures than the α subunits. The β subunits are responsible for recognizing nucleotide binding and dissociation at the catalytic sites, which facilitate the open-to-closed motions (4, 5). The γ subunit undergoes a 120° rotational step, accompanying the hydrolysis of one ATP molecule, as demonstrated by single-molecule rotation measurements using *Bacillus* PS3 F_1_-ATPase (TF_1_) (6). Furthermore, single-molecule rotation measurements under low ATP concentrations close to *K*_m_ elucidated that a 120° step of TF_1_ can be divided into an 80° substep associated with ATP binding and a 40° substep linked to ATP hydrolysis (7, 8) (Fig. S1). The pauses preceding each substep have been referred to as the binding dwell state and catalytic dwell state, respectively (8, 9). Interestingly, F_1_-ATPases from other species have different substep rotation mechanisms. For instance, human mitochondrial F_1_ (hMF_1_) and bovine mitochondrial F_1_ (bMF_1_) have 65° + 25° + 30° substep rotations (10) and 10∼20° + 70∼60° + 40° substep rotations (11), respectively.

**Figure 1.**
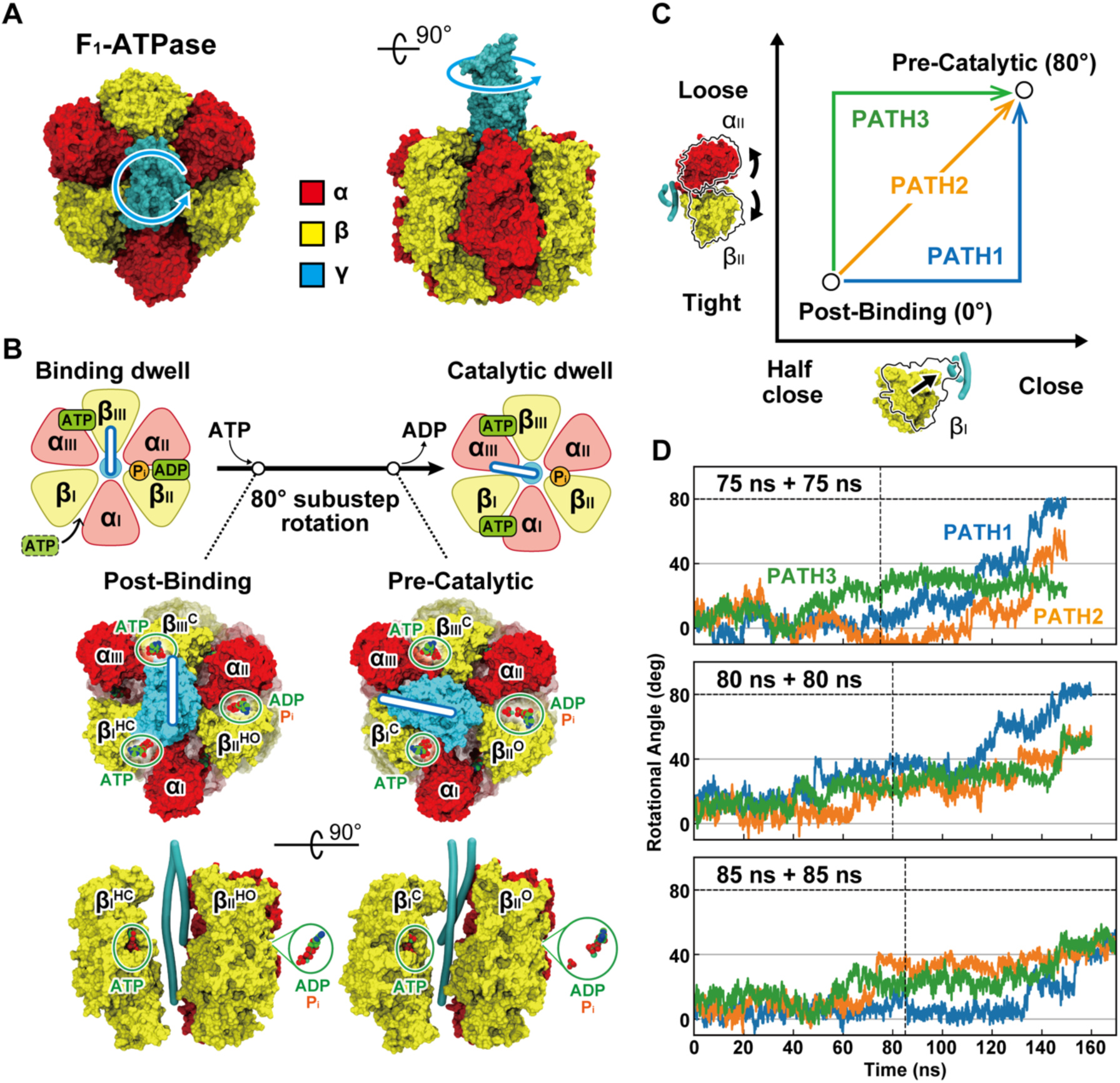
Structure of F_1_-ATPase and angular rotation of γ subunit by Targeted MD. A: F_1_-ATPase viewed from the membrane side (left) and from the side (right) (PDB ID: 7L1Q). The rotor γ subunit (cyan) is embedded in the stator ring α_3_β_3_, which is formed by α subunits (red) and β-subunits (yellow). The γ subunit rotates counterclockwise during hydrolysis. B: Schematic representation of an 80° substep of TF_1_. As experimentally demonstrated, the structures indicating binding dwell and catalytic dwell are illustrated with the α subunits (red) and β subunits (yellow) represented as rounded triangles and the occupancy states of the catalytic sites indicated. The blue sticks denote the direction of γ subunit. “Post-Binding” and “Pre-Catalytic” correspond to the binding dwell after ATP binding and the catalytic dwell structure where ADP or P_i_ has not yet dissociated, respectively. The three αβ pairs are uniquely identified as α_I_β_I_, α_II_β_II_, and α_III_β_III_, and their conformations are represented as Open (O), Half-open (HO), Half-close (HC), and Close (C). C: Structural changes illustrating the hinge motion of β_I_ and the loosening at the α_II_-β_II_ interface from binding dwell to catalytic dwell, as explored through TMD. D: Changes in the rotation angle of the γ-subunit observed through TMD. The calculation time is the sum of the initial and subsequent movements, with 75 ns + 75 ns signifying that the initial movement was induced over 75 ns of TMD, followed by the subsequent movement over another 75 ns of TMD. The three-colored time-series data correspond to the color coding in Fig. 1C, with PATH1 (blue), PATH2 (orange), and PATH3 (green) representing the analysis results of angular change analyses.

The structural details of the dwell states have been predominantly elucidated through X-ray crystallography of bMF_1_ (12–15). Since the first structure (PDB ID: 1BMF), the rotational angles of the γ subunit in most of the crystal structures of bMF_1_ are found near 40° (16), suggesting that they correspond to the catalytic dwell state (17). For the widely utilized TF_1_, the crystal structures for the stator ring lacking the γ subunit (18) and the self-inhibited state where the ε subunit impedes rotation were reported (19). Recent structural studies of TF_1_ using cryo-electron microscopy (cryo-EM) determined three structures: one for the temperature-sensitive (TS) dwell state (PDB ID: 7L1Q) and two for the catalytic dwell state (PDB ID: 7L1R, 7L1S). The TS dwell state structure shows almost 0° rotation angle of the γ subunit (20) and is very similar to the binding dwell state (PDB ID: 8HHA) of TF_o_F_1_ cryo-EM structures (21). These cryo-EM structures with different rotation angles provide a better understanding of the molecular mechanism of substep rotation.

So far, molecular dynamics (MD) simulation studies of F_1_-ATPase have been extensively conducted utilizing the catalytic dwell structures of bMF_1_, E. coli F_1_, and yeast F_1_. These MD studies can be broadly divided into three approaches. The first one involves inducing structural changes in the stator and examining the resulting changes in the rotor (22–24). The second approach applies an external force to the rotor to force it to rotate and examines the resulting structural changes in the stator (25–29). The third one focuses on observing combinations of stator and rotor structures (30–32). Coarse-grained (CG) MD studies have primarily employed the first and third approaches, investigating the origins of the rotational driving force. Several research groups suggested that the steric interactions between the β and γ subunits associated with the opening and closing motions of the β subunit can drive γ-rotation using CG MD simulations (22, 23). Contrasting results were reported that electrostatic interactions between the β and γ subunits play a significant role (30, 31). All-atom MD studies have usually employed the second approach, wherein the γ subunit is enforced to rotate. They investigated the dynamic properties of the γ subunit (25, 27, 33), the timing of phosphate release (26), and the prediction of ATP binding states (28). Also, most of the all-atom MD studies utilized only the catalytic dwell structures.

In this study, we conducted all-atom MD simulations of TF_1_ to understand the molecular mechanisms underlying the 80° substep rotation. There are two key differences between the current MD study and previous ones. First, we utilized the recent cryo-EM structures that show rotation angles close to the binding dwell and catalytic dwell states, which are the two endpoint structures during the 80° substep rotation. Secondly, we utilized the first approach, wherein the conformational changes of the stator α_3_β_3_ are induced first, and spontaneous γ-rotations are examined. This approach is free from slow responses of the stator conformations upon rapid γ- rotations induced by strong external forces. For this purpose, we performed targeted MD (TMD) simulations (34) for initial transition path prediction, the mean-force string method (35) for the path optimization, and the umbrella sampling (US) (36) to obtain the potential mean force (PMF) along the optimized pathway. In the mean-force string method and the umbrella sampling simulations, 64 images of TF_1_ solvated with explicit water, which consists of more than 500,000 atoms, were simulated in parallel. We used the supercomputer Fugaku at RIKEN Center for Computational Science and MD software GENESIS (37–39) to perform the computationally expensive simulations. Based on the simulation results, we propose a new atomistic mechanism for the γ-rotation in F_1_- ATPase, which we call the distortion-push mechanism. This mechanism can explain many experimental results and answer the long-term questions about the rotational mechanisms underlying detailed mechanisms for F_1_-ATPase.

## Results

### TF_1_ structures before and after 80° substep rotation

Fig. 1B shows a schematic representation of 80° substep rotation, transitioning from the binding dwell to the catalytic dwell state. It is induced by ATP-binding at the catalytic site of one of the αβ dimers (α_I_β_I_). ADP at the catalytic site of the second αβ dimer (α_II_β_II_) is released during 80° substep rotation. ATP at the catalytic site of the third αβ dimer (α_III_β_III_) remains intact during the substep rotation and is hydrolyzed into ADP and P_i_ in the following 40° substep rotation. Sobti et al. determined one structure in the TS-dwell state, wherein the β_I_ subunit is half-closed and ATP is loosely bound at the β_I_ catalytic site (PDB ID: 7L1Q), and two structures in the catalytic dwell state: one with ADP at the interface of the α_II_β_II_ dimer (PDB ID: 7L1R) and another with P_i_ at the same interface (PDB ID: 7L1S)(20). The TS-dwell structure is very close to the binding dwell structure of TF_o_F_1_ (PDB ID: 8HHA), which was later determined with cryo-EM (21). The Cα atom root mean square deviation (RMSD) between 7L1Q and 8HHA is 0.84 Å. The binding dwell structure (PDB ID: 8HHA) was unavailable when we started this study. We, therefore, used the TS-dwell structure as the starting conformation in the 80° substep rotation. We added P_i_ to the cryo-EM structure in the catalytic dwell state bound with ADP (PDB ID: 7L1R). These two structures, which are referred to as Post-Binding and Pre-Catalytic states, include identical nucleotides at three active sites: ATP at the catalytic sites of β_I_ and β_III_; ADP and P_i_ at the interface of the α_II_β_II_ dimer. Thus, we could focus on conformational changes of the stator α_3_β_3_ and the responded γ-rotation in MD simulations.

### Transition pathways in 80° substep rotation predicted with Targeted MD

Sobti et al. found two essential conformational changes between the TS-dwell (7L1Q) and catalytic dwell (7L1R, 7L1S) structures (20): one is the hinge motion of β_I_ (from half-closed to closed), and another is the interface motion of the α_II_β_II_ dimer (from tight to loose). Previous structural studies also pointed out the importance of these motions (16, 40). We designed three possible pathways during 80° substep rotation by changing the order of the two essential motions (Fig. 1C). In PATH1, the hinge motion of β_I_ precedes the interface relaxation of the α_II_β_II_ dimer. The two motions undergo simultaneously in PATH2. PATH3 is opposite to PATH1: the interface relaxation of the α_II_β_II_ precedes the hinge motion of β_I_. To induce the conformational changes in F_1_-ATPase with TMD, external forces were applied to β_I_ and the α_II_β_II_ dimer interface.

TMD simulations from Post-Binding to Pre-Catalytic structure along PATH 1, 2, and 3 were performed over three simulation times: 150 ns, 160 ns, and 170 ns (Fig. S2). The 80° substep rotation of the γ subunit was accomplished in two TMDs along PATH1 (150 ns and 160 ns) (Fig. 1D). TMD along PATH2 and PATH3 exhibited unidirectional counterclockwise rotations but did not realize 80° rotation of the γ subunit. Interestingly, one of the TMDs along PATH1 (170 ns) did not reproduce 80° rotation, suggesting that further simulations without external forces of TMD are necessary. We used a transition-path trajectory from TMD along PATH1 (160 ns) as an initial path for the mean-force string method simulations (35), enabling a more detailed analysis on the rotational mechanism. Although not the focus of this study, ATP always remains bound at the non-catalytic interfaces (α_I_β_II_, α_II_β_III_, α_III_β_I_).

### The mean-force string method suggests two rotational steps with resting in between

In the mean-force string method simulation, Cartesian coordinates of the Cα atoms near the subunit interface in F_1_-ATPase were used as the collective variables (CVs) (41). From the TMD trajectory along PATH1 (160 ns), we selected 64 equally spaced structures in the high-dimensional CV space. Previous studies demonstrated the effectiveness of this method in elucidating the functional rotation pathway of the multidrug transporter AcrB (42) and in analyzing the E1P-E2P transition of the Sarcoplasmic Reticulum Ca²⁺-ATPase (43). By performing the string method simulations for 82 ns per image, the initial pathway was optimized to a free energy minimum pathway, with convergence evaluated based on the RMSD changes of each image (Fig. S3). Subsequently, the umbrella sampling (US) simulation was performed over 100 ns for each replica representing this transition pathway (Fig. S4). PMF was computed using the multistate Bennett acceptance ratio (MBAR) from the trajectories over the final 90 ns (Fig. 2A).

**Figure 2.**
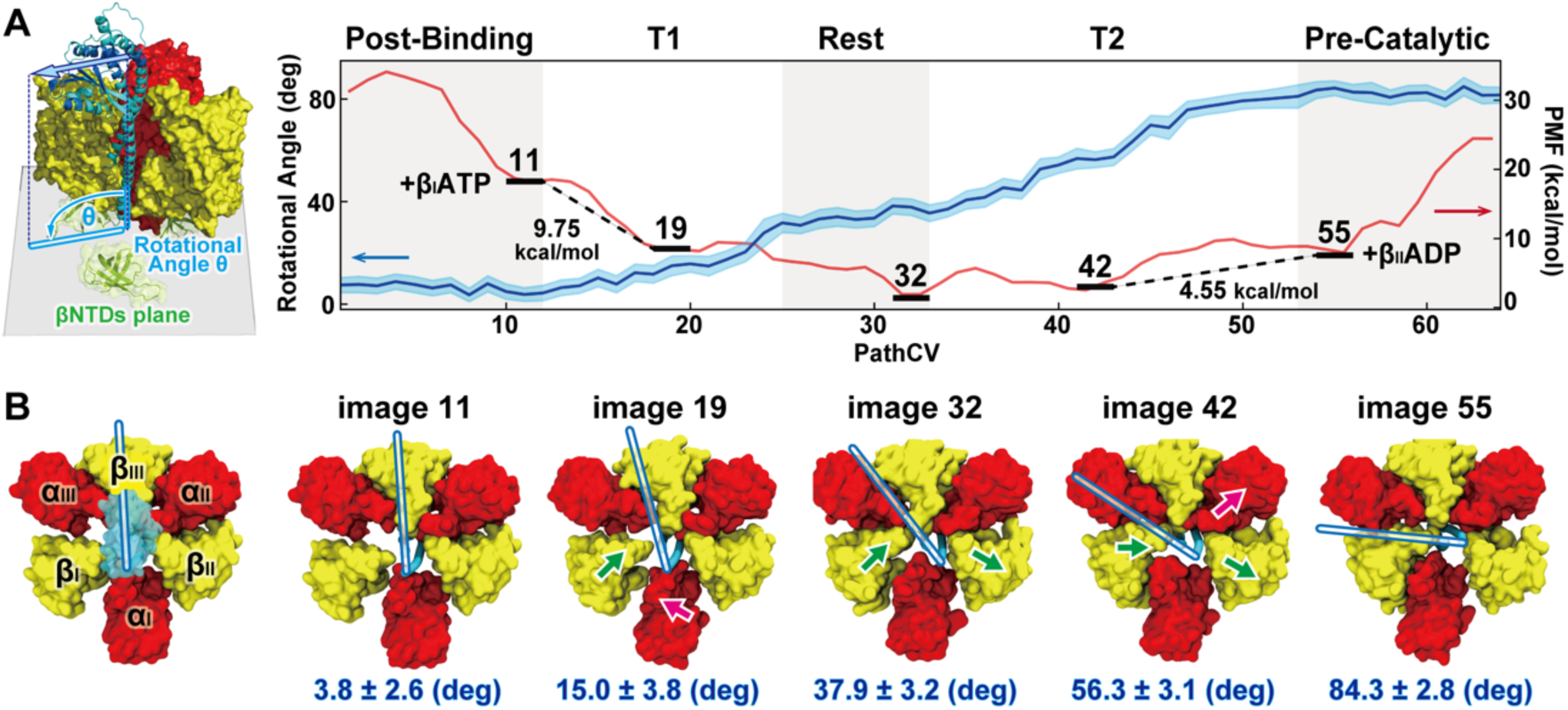
Conformational change mechanism of 80° substep rotation A: Rotational angle changes of the γ subunit predicted by the string method (blue) and the free energy profile (red). The five regions, color-coded in gray and white, represent the five phases classified based on the rotational angle changes. B: Representative structures of the five phases, using configurations obtained after 100 ns of umbrella sampling.

We calculated the average γ-rotational angles from the trajectory of each replica and obtained five different phases using the change point detection: images 1-12, 13-25, 26-33, 34-53, and 54- 64 (Fig. 2A). The phase including the image 1-12 has the average γ-rotational angle of 6.3±3.7°, suggesting that this phase is stable with a minimal change of the γ-rotation. Similarly, the phase, including images 54-64, shows stable rotation angles with the highest average values (82.4±3.5°). Thus, we refer to the first and 5^th^ phases as Post-Binding and Pre-Catalytic states, respectively. Significant changes in the average γ-rotational angle were observed in the second (images 13-25: from 6.3° to 20.4°) and fourth phases (images 34-53: from 37.1° to 79.8°). We refer to them as T1 and T2 steps, respectively. Between T1 and T2, the γ-rotational angles do not change significantly. This “Rest” phase covers the images 26-33 (34.5±4.1°). Two rotation steps (T1 and T2) and the Rest in between can characterize the 80° substep rotation from Post-Binding to Pre-Catalytic state. Fig. 2A shows PMF along the minimum free-energy pathway obtained with the string method.

The metastable states that represent Post-Binding, T1, Rest, T2, and Pre-Catalytic states are identified in images 11, 19, 32, 42, and 55, which reveal the γ-rotational angles of 3.8±2.6°, 15.0±3.8°, 37.9±3.2°, 56.3±3.1°, and 84.3±2.8°, respectively (Fig. 2B). The PMF in Fig. 2A is characterized by elevated free energy at both ends, suggesting that a large free-energy difference between the binding dwell and Post-Binding, and that between the catalytic dwell and Pre-Catalytic state. Since the structural changes between them and the γ-rotational angle differences are small, such significant free-energy changes can be due to the differences in the nucleotide-bound states. To investigate this possibility, we performed two additional US simulations. In one US simulation from images 1 to 32, we removed ATP from the β_I_ active site, creating structures with the same substrate-bound state as the binding dwell state (Fig. S5). As a result, image 11 (the binding dwell state) is 8.45 kcal/mol more stable than image 19 (T1) (Table S1). The ATP binding at the β_I_ active site can create a downhill free-energy landscape to drive the initial rotation of the γ subunit (Fig. 2A). In the second US simulation, ADP at the interface of the α_II_β_II_ dimer was removed to mimic the catalytic dwell structure. In this state, P_i_ remained bound to the catalytic site. We used images 33- 64 in the US and found that image 55 is stabilized by 1.60 kcal/mol compared to image 42 (Table S1). This result suggests the catalytic dwell structure is stabilized after ADP is released from the Pre-Catalytic state. By integrating the three US results, the reaction model from the binding dwell to the catalytic dwell state agrees with that proposed by single-molecule measurements (44).

### The stator-ring conformational changes during 80° substep rotation

Fig. 3A suggests that the hinge motion of β_I_ starts from the half-closed state in T1 and continues toward the closed state in Rest and T2. At the same time, the α_II_β_II_ interface relaxation remains unchanged before T2 (Right panel of Fig. 3B and Fig.S6). This order of two essential motions is the same as PATH1 in TMD, while T2 includes both. The hinge motion of β_II_ happens from half-open to open form in T2, synchronizing the interface relaxation of the α_II_β_II_ dimer (Fig. S6). The α_I_β_I_ interface becomes tighter during 80° substep rotation. The α_III_β_I_ interface completes tightening by image 32 in Rest, remaining stable since then (Left panel of Fig. 3B).

**Figure 3.**
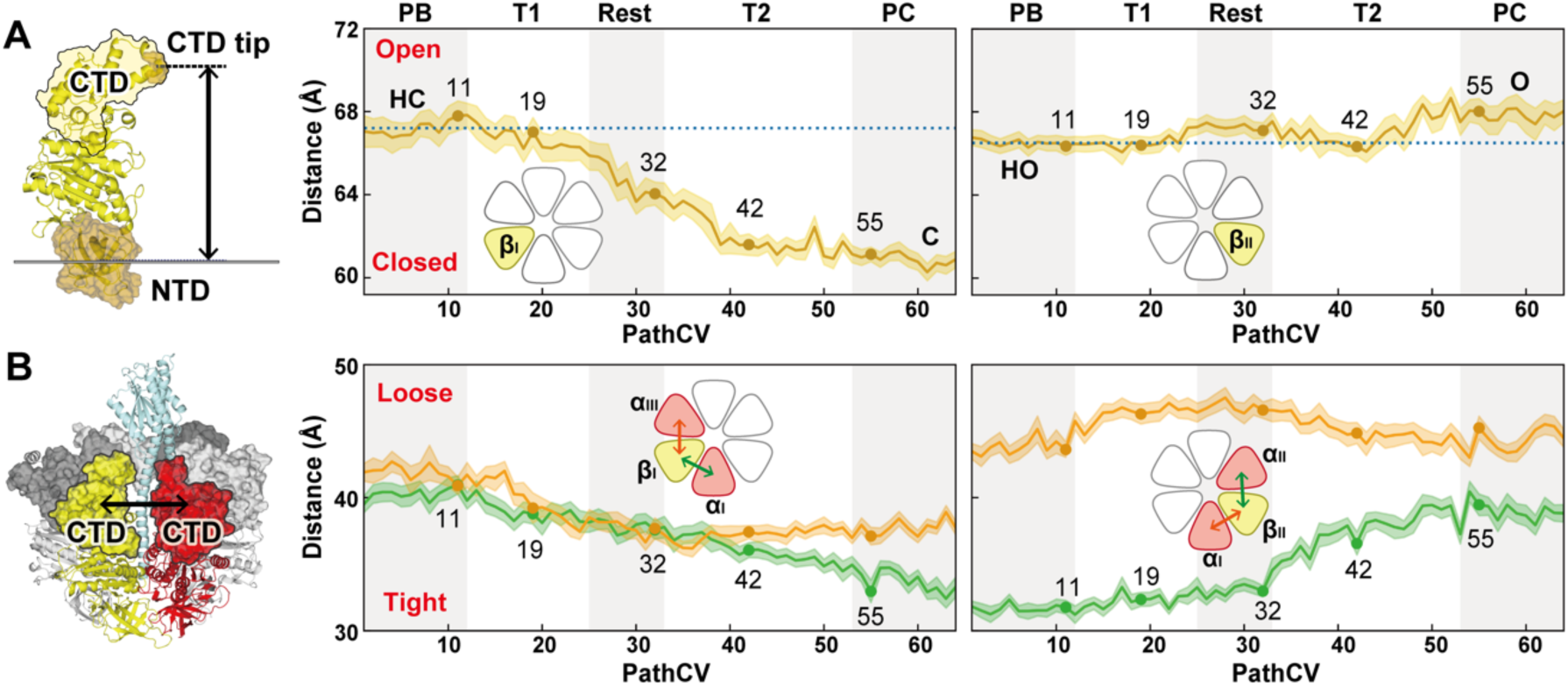
Domain motion of the stator during 80° substep A: Distance between the N-terminal domain (NTD) and the C-terminal domain (CTD) tip representing the hinge motion of β_I_. B: Distance between the CTD of the α and β subunits, indicating the relaxation at the α-β interface.

### Interaction between γ-rotator and α_3_β_3_ stator to drive 80° substep rotation

The stator ring and γ subunit make contacts at the top (orifice region) and bottom (switch II region and hydrophobic sleeve region) in the X-ray structures (12). As the orifice region containing the C-terminal domain (CTD) of the β subunit is known to be responsible for at least half of the torque generation (45, 46), we focus on interaction changes in the region. In T1, residues in the orifice region of β_I_ interact weakly with Q183 in the γ subunit (Fig. 4A), and there are no significant interaction changes between the γ and β_I_ subunits at the orifice and switch II regions (Fig. S7, S8). Instead, the electrostatic interaction changes between the γ subunit and the stator opposite β_I_ are essential in T1 (Fig. 4B, S9, S10). At first, R88 and G89 in the loop region of the γ subunit change their interaction partners from β_II_E391 to β_III_E391 and from α_II_D401 to β_II_E391, respectively. In addition, a hydrogen bond between the backbone of γL90 and the sidechain of β_III_E391 is broken. The interaction changes associated with the start of T1 also occur within the stator ring: the three interfaces of α_I_β_I_, α_III_β_III_, and α_II_β_II_ form new interactions by switching their interaction partners (Fig. S11). Thus, T1, the 35° rotation in 80° substep rotation, is not driven by the direct interaction between the due to the distortion of the stator ring without direct contact between β_I_ and γ subunits, but the distortion of the whole stator ring and the electrostatic interactions between the γ subunits and the stator opposite β_I_ are the two key driving forces.

**Figure 4.**
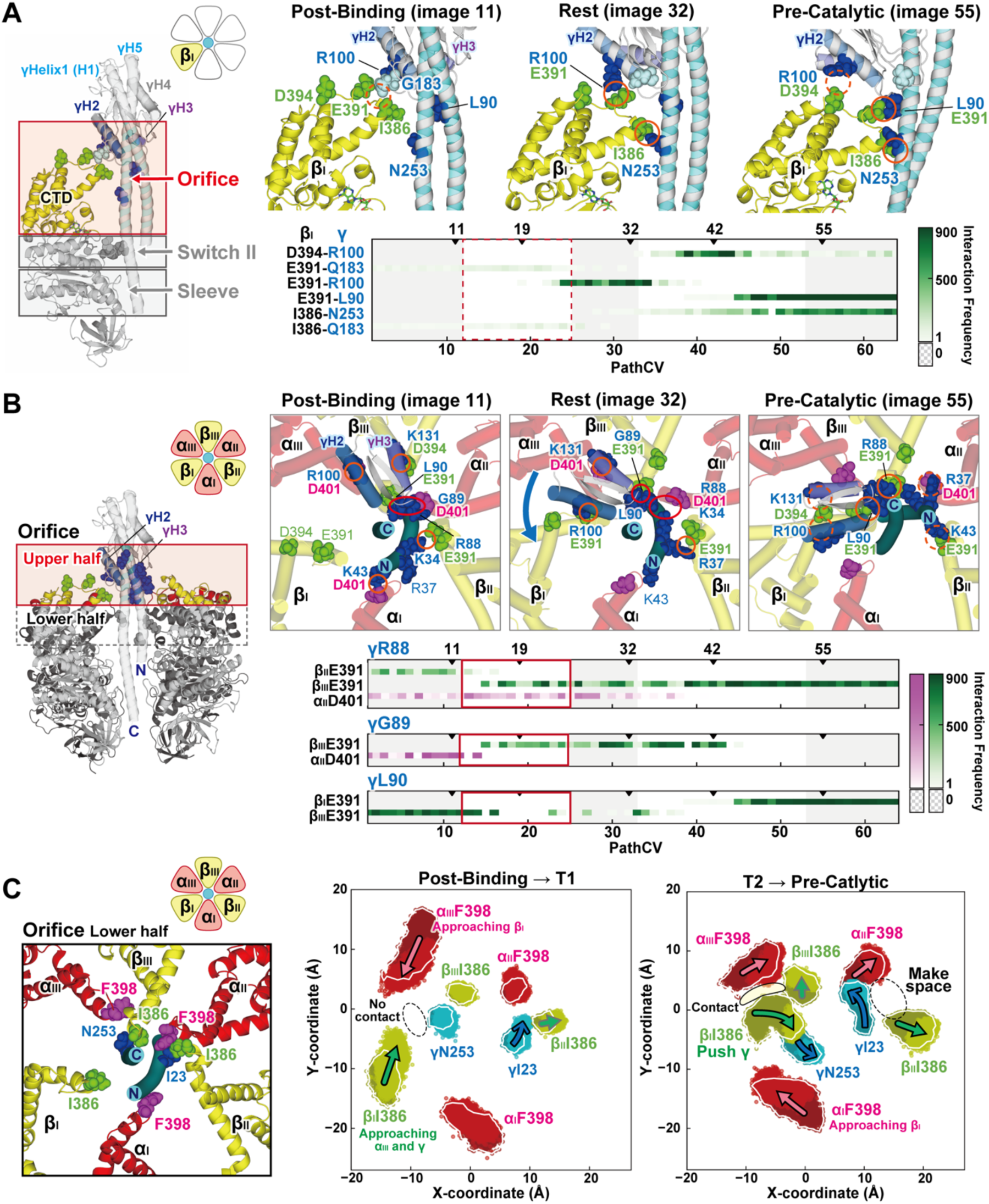
Contacts changes between the α_3_β_3_ and the γ subunit A: Changes in the interaction between the CTD of the β_I_ subunit and the γ subunit. Residues in the β_I_ and γ subunits exhibiting notable interaction changes during the 80° rotation are highlighted as green and blue spheres in images 11, 32, and 55. The weakly interacting γG183 is shown as a light blue sphere. The graph below illustrates the interaction frequency for each residue pair, with higher frequencies observed during 900 frames of the 90 ns umbrella sampling indicated by darker colors. Interaction changes in the switch II between the β_I_ and γ subunits are provided in Fig. S8. B: Interaction changes between the upper CTD region of the α_3_β_3_ stator ring and the γ subunit are shown. Residues in the α, β, and γ subunits that form significant interactions between the upper CTD region of α_3_β_3_ and the γ subunit are highlighted as magenta, green, and blue spheres, respectively, in images 11, 32, and 55. The graph below represents the interaction changes involving γ88-90 and α_3_β_3_. Further details on the interaction changes in the upper CTD region (excluding these residues) and the lower CTD region are available in the SI Appendix (Fig. S9 and S10, respectively). C: The trajectories of residues located at the subunit interfaces during the 80° rotation. The positional changes of αF398 and βI386 at the CTD termini of the stator ring, as well as those of γ subunit residues I23 (N-terminal helix) and N253 (C-terminal helix) in contact with these residues, are plotted on a plane as viewed from the top of TF_1_. The trajectories show transitions from Post-Binding and T2 states (darker colors) to T1 and Pre-Catalytic (brighter colors).

At the end of T1, just before Rest (image 24), β_I_E391 interacts with R100 in Helix2 (γH2). This electrostatic interaction switches those between β_I_D394 and γR100 in T2, which, in turn, shifts into those between β_I_E391 and γL90. In addition, I386 at the CTD tip of β_I_ gradually strengthens the interaction with N253 in the γ-coiled coil, suggesting that T2, the second rotation after Rest, is driven by direct interactions between β_I_ and γ subunits.

This model is confirmed by the contact analysis between the stator ring and the γ subunit at the lower half of the orifice region (Fig. 4C). In the projection viewed from the top, β_I_I386 moves towards α_III_F398 and γN253, and α_III_F398 approaches β_I_I386 from Post-Binding to T1. At the end of T1, β_I_I386 and γN253 are still separated by about 5 Å (Fig. 4C, S12). From T2 to Pre-Catalytic phase β_II_I386 moves away from the γ subunit, and the γI23 position rotates counterclockwise. β_I_I386 approaches γN253, moved to push γN253 counterclockwise. α_III_F398 likely supports the motions around β_I_I386. In this phase, α_II_F398 and β_II_I386 moved away from each other because of the loosening of the α_II_β_II_ interface (Fig. 3B, Fig. S12 and S13). Thus, we found that atomic mechanisms for 80° substep rotation can be explained with the distortion of the stator ring and the electrostatic interaction changes in the first 35° rotation, and the direct pushing force from β_I_ to γ subunit in the last 55° rotation. Hereafter, we call this two-step rotation mechanism the distortion-push Mechanism and relate this mechanism with the previous experimental observation in Discussion section.

### Roles of ATP binding and ADP/P_i_ release in 80° rotation

In Post-Binding, ATP, which is loosely bound to β_I_, has its phosphate group wrapped in the P-loop (_158_GxxGxGK[T/S]_165_) during T1, forming a specific interaction with G161, V162 and G163, forming a particular interaction (Fig. 5A). The main chain dihedral angles of G161 and G163 changed significantly to create this interaction (Fig. S14), confirming that P-loop utilizes the unique flexibility of Gly residues in the nucleotide binding. This interaction remains stable up to Pre-Catalytic. Previous studies have reported increased binding affinity between β_I_ and ATP as the major torque-generating event at the 80° substep (47). The tight binding of β_I_ and ATP at T1 can induce the β_I_ hinge motions and facilitate the γ rotation.

**Figure 5.**
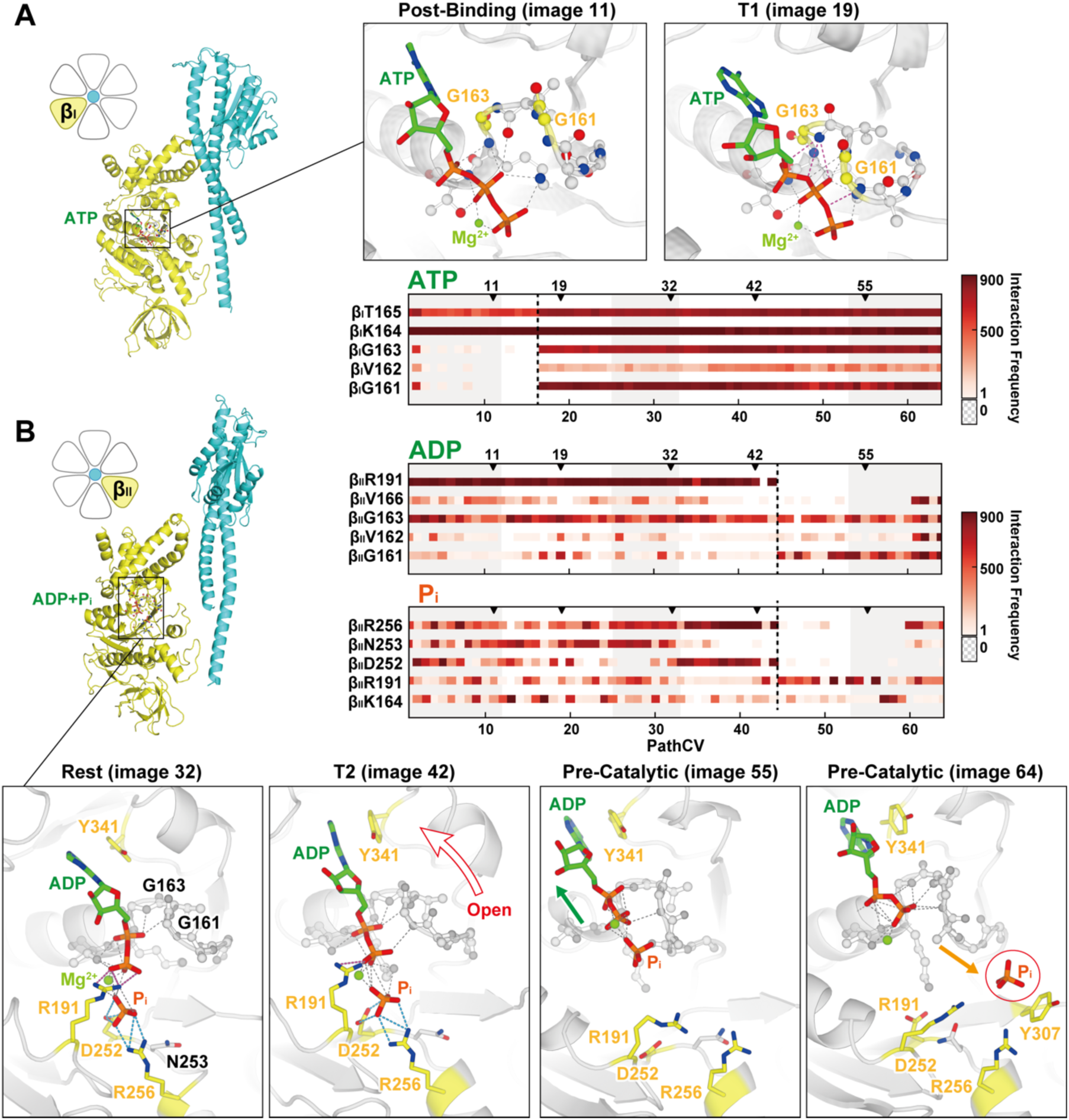
Interaction changes at the catalytic site of the β subunit A: The structure of ATP bound to the catalytic site of β_I_. During the transition from the Post-Binding state to T1, interactions between the phosphate groups in ATP and the P-loop residues G161, V162, and G163 are formed. The time-dependent changes in the β_I_ATP interactions are represented by brightness, with higher brightness indicating more frequent interactions observed during the 90 ns of umbrella sampling. B: The structure of ADP and P_i_ bound to the catalytic site of β_II_.

ADP and P_i_ bound at the α_II_β_II_ interface reduce their interactions with catalytic residues β_II_R191, D252, and R256 in T2. Then ADP and P_i_ move upwards of β_II_ with the CTD side of β_II_ (Fig. 5B). This movement coincides with the transition of β_II_ from the half-open to open state (Fig. 3A). In Rest, where these interactions remain intact, it transiently causes transitions from the half-open to open conformation before reverting. It returns to the half-open state again at the beginning of T2. These results suggest that cleavage of the interaction between ADP and β_II_R191, D252, and R256 is significant for β_II_ to maintain the open conformation. Furthermore, after T2 rotation, ADP lost its interaction with α_II_R365 and no longer interacted closely with α_II_ (Fig. S15). This implies that ADP, which does not contribute to the tightening of the α_II_β_II_ interface during Pre-Catalytic, is bound to β_II_ and can be rotated 80° by the loss of interaction with α_II_ without complete release from β_II_.

## Discussion and Conclusions

Here, we discuss the relationship between the proposed distortion-push mechanism and previous experimental/computational studies of F_1_-ATPase. Single-molecule rotation experiments have shown that the 80° substep rotation of TF_1_ is driven by the binding of ATP to β_I_ as a trigger, followed by ADP dissociation from α_II_β_II_ (44, 48). Although the present study did not directly simulate the nucleotide binding and dissociation, there was a clear trend towards an increase in the interaction between β_I_ and ATP in T1 and a decrease in the interaction between α_II_β_II_ and ADP in T2. This order is consistent with the experimentally demonstrated binding-dissociation sequence. Furthermore, in T2, the loosening of the α_II_β_II_ interface occurs after the contact formation between the β_I_ and γ subunits in Rest. These structural changes are consistent with the previous simulation results (24), which suggested that the propagation of structural changes due to β_I_ closure promotes α_II_β_II_ interface loosening and ADP dissociation. Theoretical considerations based on rotation experiments (49) also suggested the presence of a short-lived state prior to the ADP dissociation at a rotation angle of 30° during 80° substep rotation of TF_1_. Similar results have been confirmed in E.coli F_1_ rotation experiments (50) and simulations of synthetic rotation of bMF_1_ (32). The rest around 35° can be interpreted as a rotational pause waiting to loosen the α_II_β_II_ interface and the ADP dissociation.

In the cryo-EM structure (PDB ID: 7L1R), which shows the catalytic dwell state, only ADP is found to be bound to the α_II_β_II_ interface. This structure cannot explain the conventional rotational catalysis scheme in which ADP already dissociates during the 80° substep (44). The authors (20) suggested that this ADP can dissociate once and then rejoin at the same binding site. In the present study, at the end of T2, ADP reduces most of the interaction with α_II_ and loses the role in tightening the α_II_β_II_ interface. After the interactions between ADP and β_II_R191, D252, and R256 are cleaved in T2, the closing of the β_II_ hinge cannot happen. Therefore, this study suggests that a complete dissociation of ADP from β_II_ is not required for the catalytic dwell state after 80° rotation and that ADP, which is not involved in driving the rotation of the γ subunit, can rebind to β_II_. Recent cryo-EM structures under high ATP concentrations have shown that ATP can bind to α_E_β_E_ (α_II_β_II_ around 80° rotation) independent of rotation angle (21). The role of bound nucleotide at the β subunit in the rotation mechanism should be discussed carefully together with the F_1_-ATPase structure and the detailed interaction modes.

The most dominant rotational driving force of the γ subunit rotation has been considered to be either electrostatic interaction (30, 31, 41) or steric hindrance (22, 23, 51). The distortion-push mechanism proposed here can answer the question, suggesting different driving forces for the two rotational steps intervening by Rest. In this mechanism, the first rotation is driven by the distortion of the stator and the electrostatic interaction, while the direct contact between the β and γ subunits. Therefore, this molecular mechanism can be considered as one that encompasses two different models. It also agrees with the previous free-energy calculations on the ATP synthesis rotation of bMF_1_ (32), which suggested that the rotation is mediated by a more complex combination of electrostatic and steric interactions in the rotation after ATP dissociation.

In summary, the molecular mechanism underlying 80° substep rotation of F_1_- ATPase (TF_1_) from the thermophilic bacterium *Bacillus* PS3 was investigated using all-atom MD calculations. The optimized transition path and PMF along the path revealed that 80° substep rotation consists of two rotations separated by a rest around 35°. The distortion of the stator α_3_β_3_ and the electrostatic interaction changes between the γ subunit and αIIβII or βIII subunits in the upper part of the orifice region drive the first rotation. The γ rotational angle does not change significantly in Rest, while β_I_ continues the hinge-bending motion and approaches to form a tight interface with α_III_. The CTD of β_I_ makes contact with the coiled-coil part of the γ subunit. In the second rotation, the CTD of β_I_ pushes the coiled-coil region of the γ subunit by steric repulsion. The interaction between the CTD of α_II_β_II_ is broken. The α_II_β_II_ interface gradually starts to loosen, and thereby ADP reduces interaction with the catalytic residues of β_II_ (R191, D252, R256) and β_II_ transitions to an open conformation. As we see in the current study, chemo-mechanical coupling in F_1_-ATPase is explained with multiple factors, such as distortion of the stator α_3_β_3_ subunits, changes of the electrostatic interaction between the β and γ subunit, the strengths of nucleotide-binding at each αβ interface, and so on. Proper molecular modeling and simulations based on the cryo-EM structures could help our understanding of atomistic mechanisms for the substep rotation of the γ subunit. The current approach can apply to the following 40° substep rotation toward a complete understanding of an entire rotational cycle in F_1_-ATPase.

## Materials and Methods

### Computational setup for modeling and simulations of TF_1_

In this study, the binding dwell and catalytic dwell states of TF_1_ were taken from the cryo-EM structures corresponding to the rotational angles of each state (PDB ID: 7L1Q and 7L1R, respectively). The protonation state of P_i_ at the ADP+P_i_ binding site of each state was set to a doubly protonated state (H_2_PO_4−_), as suggested by previous QM/MM studies (5, 52, 53). Furthermore, αE165 and αE272 were assigned to protonated form based on the pKa predictions from PROPKA (54, 55). The internal cavities of the protein were hydrated using Dowser++ (56), and the structures were modeled using the VMD plugin (57). Each structure was solvated in a cubic box with a side length of 174.8 Å, containing CHARMM TIP3P water molecules (58) and neutralized with 150 mM KCl. The total atom count of the solvated systems was approximately 513,000 atoms. The CHARMM c36m force-field parameters (59) were used for TF_1,_ and the refined parameters for nucleotides by Komuro et al. were employed (60). Water molecules are considered as rigid using SETTLE constraints (61). All bonds involving hydrogen atoms are constraints with SHAKE/RATTLE (62, 63). All molecular dynamics (MD) simulations were performed using the GENESIS software version 2.1 (37–39). The simulations were conducted in the NPT ensemble at 310 K and 1 atm, using the stochastic velocity rescaling thermostat and MTK style barostat (64, 65) using the refined temperature definition (66). Group temperature/pressure is used for evaluation of temperature/pressure in thermostat/barostat (67, 68). The nonbonded interactions were calculated using the CHARMM standard procedures: the LJ interaction was smoothly truncated from 10 Å to 12 Å using the force-based switch function (69), and the electrostatic interactions were computed with the particle mesh Ewald summation algorithm (70, 71). For all executions, except for equilibration, we utilized the r-RESPA (or multiple time step) integration with 2.5 and 5.0 fs time steps for the real- and reciprocal-space interactions, respectively (72).

### Transition pathways from the binding dwell to the catalytic dwell state

Using targeted molecular dynamics (TMD) (34), we generated candidates of transition pathways from the binding dwell state to the catalytic dwell state by inducing the conformational changes in β_I_, α_II_, and β_II_, which are involved in the TF_1_ substrate binding and release. As shown in Fig. 1C, three distinct pathways (PATH1, 2, and 3), assuming sequential or simultaneous conformational changes in β_I_ and α_II_β_II_, were examined in TMD simulations. TMD simulations along with each pathway were performed three times on different time scales, 150 ns, 160 ns, and 170 ns. We selected a conformational transition predicted in a 160 ns TMD, assuming PATH1 as the initial pathway for the string method, comprising 64 images. The collective variables (CVs) were defined following the previous study for V_1_-ATPase (41), where the CVs were specified by the Cartesian coordinates of the Cα atoms of the residues within 5 Å of the inter-subunit interfaces of all the α, β, and γ subunits, as well as the heavy atoms of the substrates MgATP and MgADP, excluding P_i_ and the Cα atoms in the CTD of all the β subunits. The transition pathway was initially equilibrated so that 64 images were evenly positioned in the CVs space. Each image was subjected to MD simulation with a positional restraint of 0.01 kcal/mol/Å^2^ for 1 ns per image. Subsequently, the restraint strength was increased to 1.0 kcal/mol/Å^2^, and each image was equilibrated further with MD simulations of 50 ns per image. After equilibration, the transition pathway was refined using the string method under positional restraint of 1.0 kcal/mol/Å^2^, with MD simulations of 20 ns per image. During the refinement process, all images except for the endpoints were updated. Finally, MD simulations based on the string method with 64 images were carried out for 10 ns per image with a 0.1 kcal/mol/Å^2^ positional restraint to obtain the converged minimum free-energy pathway without fixing the endpoint images.

### Free-energy profiles from umbrella sampling

We performed umbrella sampling simulations with 64 replicas along the minimum free-energy pathway, followed by the string-method simulations. The collective variables (CVs) used in the umbrella sampling were the exact Cartesian coordinates of the atoms, the same as those in the string-method simulations. The CVs were restrained using a force constant of 0.02 kcal/mol/Å^2^.

Each umbrella window was simulated for 100 ns in the NPT MD simulations. The free-energy profile was analyzed using the MBAR (Multistate Bennett Acceptance Ratio) method (73), using the 90 ns of sampling data, excluding the initial 10 ns trajectories.

### Rotational angles of the γ subunit

To define the rotational angle of the γ subunit, we followed the viscoelastic model for the segment 3 of MF_1_ by Okazaki et al. (33)(Fig. 3A). First, we compute a vector connecting the center of mass (COM) of the Cα atoms of the entire region to the COM of the N-terminal helix (residues: 44-53) in the γ subunit with the highly rigid region (residues: 44-53, 78-82, 105-119, 134-139, 148-160, 162, 168, 170-174, 188-192, 222-231). This vector is projected onto the plane formed by the N-terminal domains (NTD, residues: 2-82) of the three β subunits. The projected vector can mimic the rotation vector measured in single-molecule rotational measurements and is referred to as the rotating vector. The rotational angle was computed by taking the inner product of the rotational vector of the reference structure (PDB ID: 7L1Q), which has the same rotation angle of the binding dwell state.

### Hinge motions of the β subunit and tightening/relaxation motions at the α-β interface

The hinge motion of the β subunit is quantified using the inter-domain vector between the COMs of the NTD and the CTD tips of Cα atoms (residues: 385-388). This height component is quantified using the perpendicular component of the inter-domain vector to the plane formed by the NTD of the three β subunits. The tightening and relaxation at the αβ interface are quantified using the distance between the COMs of the Cα atoms in the CTD of the α and β subunits (residue: 371-499 and 360-473).

## Acknowledgments

We used the computer resources provided by HPCI system research project (Project ID: ra000003, ra240003, hp230111, and hp240047) and by RIKEN Advanced Center for Computing and Communication (for HOKUSAI BigWaterfall, project Q22535, Q22536, and Q23536) to perform molecular dynamics simulations in this study. One of the authors (M.M.) is supported by RIKEN Junior Research Associate (JRA) Program, and M.O. was supported by RIKEN Special Postdoctoral Researchers (SPDR) Program. This work was supported in part by RIKEN pioneering projects “Biology of Intracellular Environments” and “Glycolipidologue Initiative” (to Y.S.), RIKEN incentive fund (to J.J), MEXT JSPS Kakenhi (grant number 19H05645, 21H05249 (to Y.S.), 21H05282 (to J.J)), MEXT program for Big-data-driven bio/synthetic polymer science to create absolutely circular materials (JPMXP1122714694) and Data-Driven Research Methods Development and Materials Innovation Led by Computational Materials Science (JPMXP1020230327) (to Y.S.), PRESTO grant from Japan Science and Technology Agency (JPMJPR22E2) (to M.O.).

## Notes

### Competing Interest Statement

The authors have declared no competing interest.

