## Supporting Information for "The distortion-push mechanism for the γ-subunit rotation in F1-ATPase"

Masahiro Motohashi[1,2], Mao Oide[2,5], Chigusa Kobayashi[3], Jaewoon Jung[2,3], Eiro Muneyuki[1], Yuji Sugita[2,3,4,*]
[1]Department of Physics, Faculty of Science and Engineering, Chuo University, Tokyo 112-8551, Japan.
[2]Theoretical Molecular Science Laboratory, RIKEN Cluster for Pioneering Research, Saitama 351-0198, Japan.
[3]Computational Biophysics Research Team, RIKEN Center for Computational Science, Kobe 650-0047, Japan.
[4]Laboratory for Biomolecular Function Simulation, RIKEN Center for Biosystems Dynamics Research, Kobe 650-0047, Japan.
[5]Institute for Protein Research, Osaka University, Osaka 565-0871, Japan

*Corresponding to Yuji Sugita

**This PDF file includes:**

Figures S1 to S15
Table S1
Legend for Movie S1
SI Reference

**Other supporting materials for this manuscript include the following:**

Movie S1

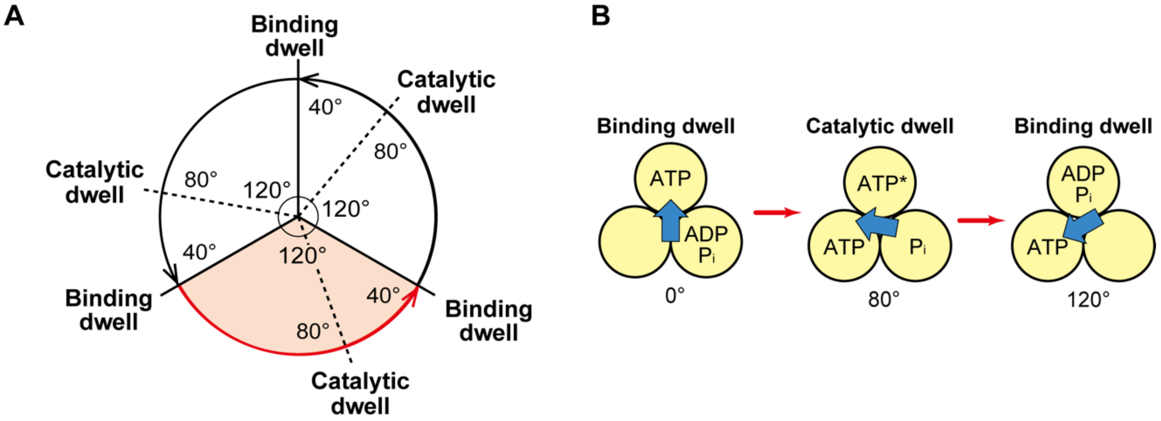


**Fig. S1. Reaction scheme of thermophilic *Bacillus* PS3 $F_1$-ATPase ($TF_1$)**

A: The 360° rotation scheme. This circular diagram illustrates a full 360° rotation consisting of three 120° steps. Each 120° step comprises 80° and 40° substep rotations. The intermediate state before the 80° rotation represents the binding dwell, while the intermediate state before the 40° rotation represents the catalytic dwell. B: The 120° rotation scheme. Yellow circles represent β subunits, with the occupancy of each catalytic site indicated within the circles. Cyan arrows show the direction of the γ subunit. During the 80° substep rotation from the binding dwell to the catalytic dwell, one ATP molecule binds to a catalytic site, and one ADP molecule dissociates from another. In the 40° substep, ATP hydrolysis and $P_i$ release occur.

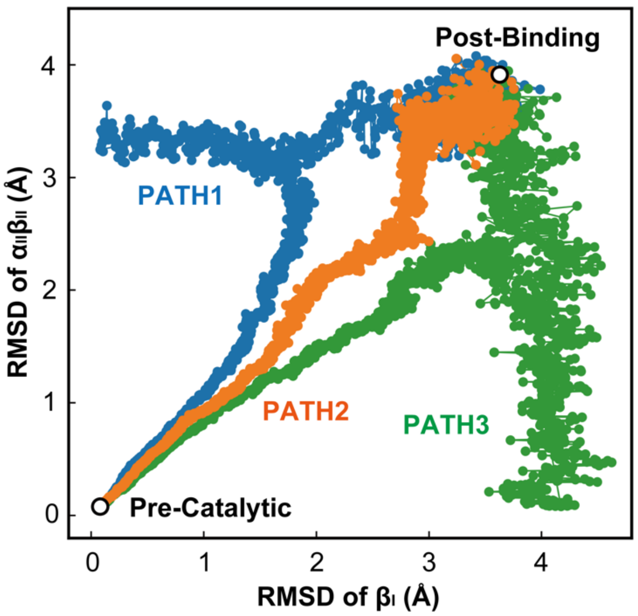


**Fig. S2. Conformational transition pathways generated by Targeted MD (TMD)**

The three pathways generated by 160 ns TMD simulations from the Post-Binding to the Pre-Catalytic state are plotted on a 2D RMSD map at 100 ps intervals. The line colors correspond to the color coding in Fig. 1C, with PATH1 (blue), PATH2 (orange), and PATH3 (green).

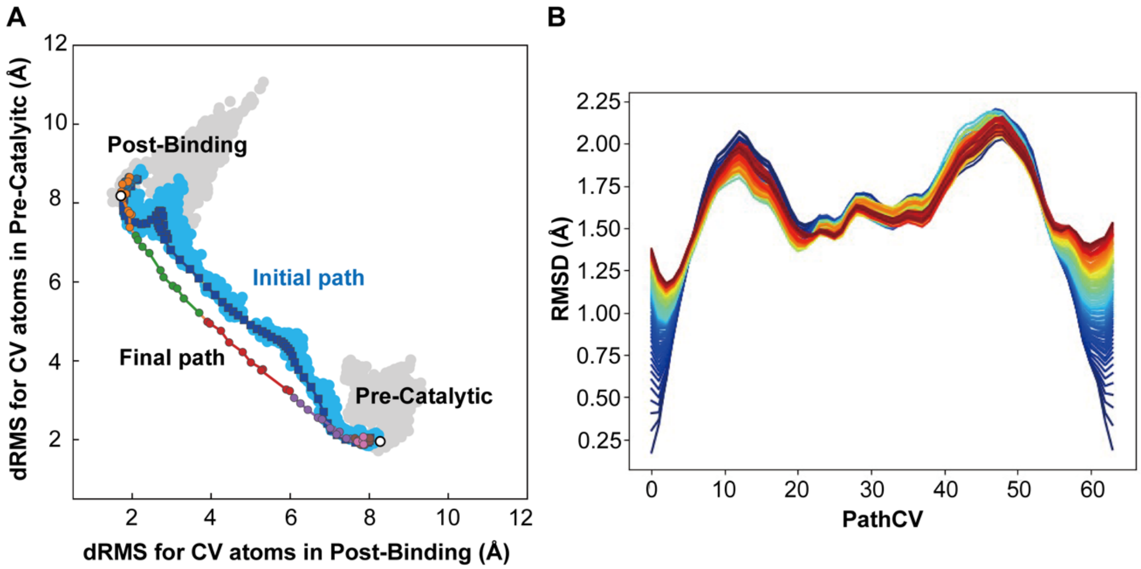


**Fig. S3. Conformational change pathway modified by mean-force string method**

A: The conformational change pathway from Post-Binding to Pre-Catalytic states. The structures obtained simulations are projected onto distance RMSDs (dRMS) from the initial and final structures, showing the trajectory of conventional MD. The initial TMD pathway is depicted in light blue and represented by 64 images in blue. This pathway was refined into the Final path using the string method. The Final path is illustrated with seven colors: blue (images 1-10), orange (11-20), green (21-30), red (31-40), purple (41-50), pink (51-60), and brown (61-64). Gray dots represent the trajectory of the conventional MD simulations from the initial to final structures. B: The convergence of the refined conformational change pathways. The colored lines represent RMSD changes between structures on the initial pathway and those on the updated pathways. Updated structures were collected every 100 ps, and the RMSD was calculated for each one and colored accordingly. The blue line represents the structures after applying the string method with a position restraint of 1.0 kcal/mol/Å² for 20 ns, while the red line shows structures following further refinement with a 0.1 kcal/mol/Å² restraint.

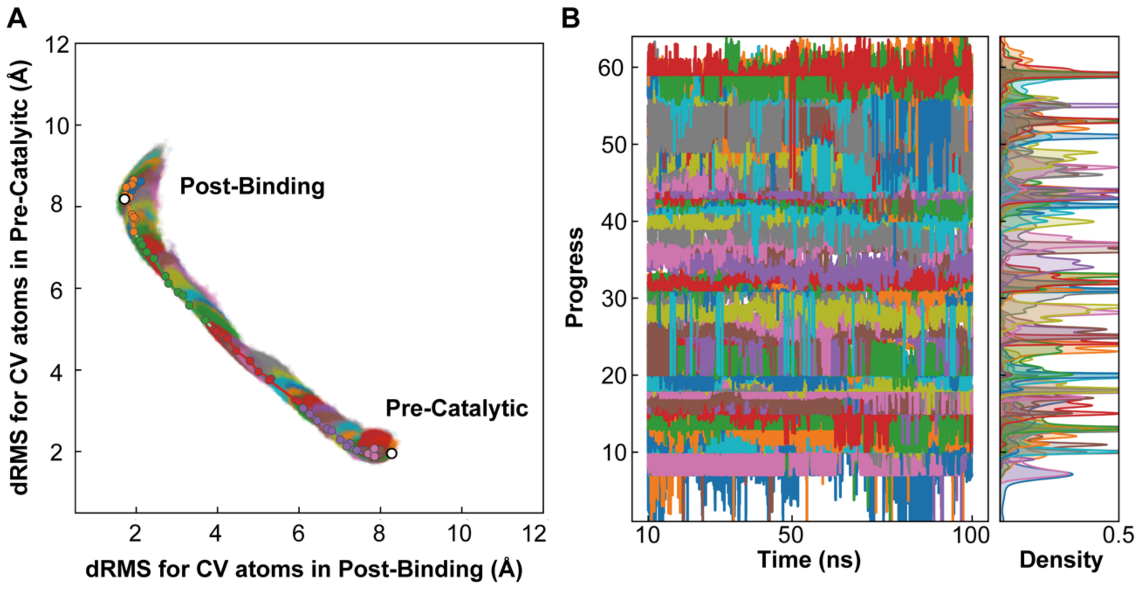


**Fig. S4. Conformational sampling on the optimized pathway using umbrella sampling.**

A: Spatial distribution along the dRMSs sampled during the last 90 ns of the 100 ns umbrella sampling (US) simulation. Dots of a distinct color represent each of the 64 sampling windows. B: Overlap of spatial sampling regions across the 64 windows.

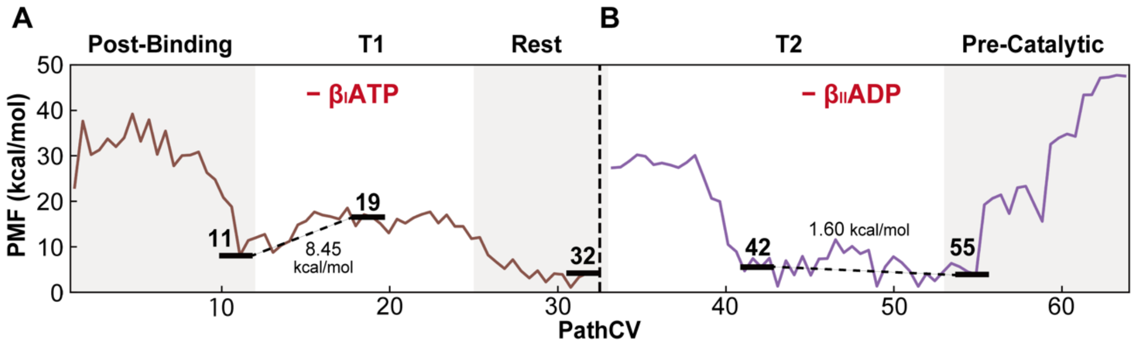


**Fig. S5. Free energy landscapes without nucleotides**

A: Brown line represents the free-energy landscape using structures from images 1 to 32, where ATP was removed from $\beta_I$. B: Purple represents the free-energy landscape using structures from images 33 to 64, where ADP was removed from $\beta_{II}$. Images 11, 19, 32, 42, and 55 represent the five major phases' structures.

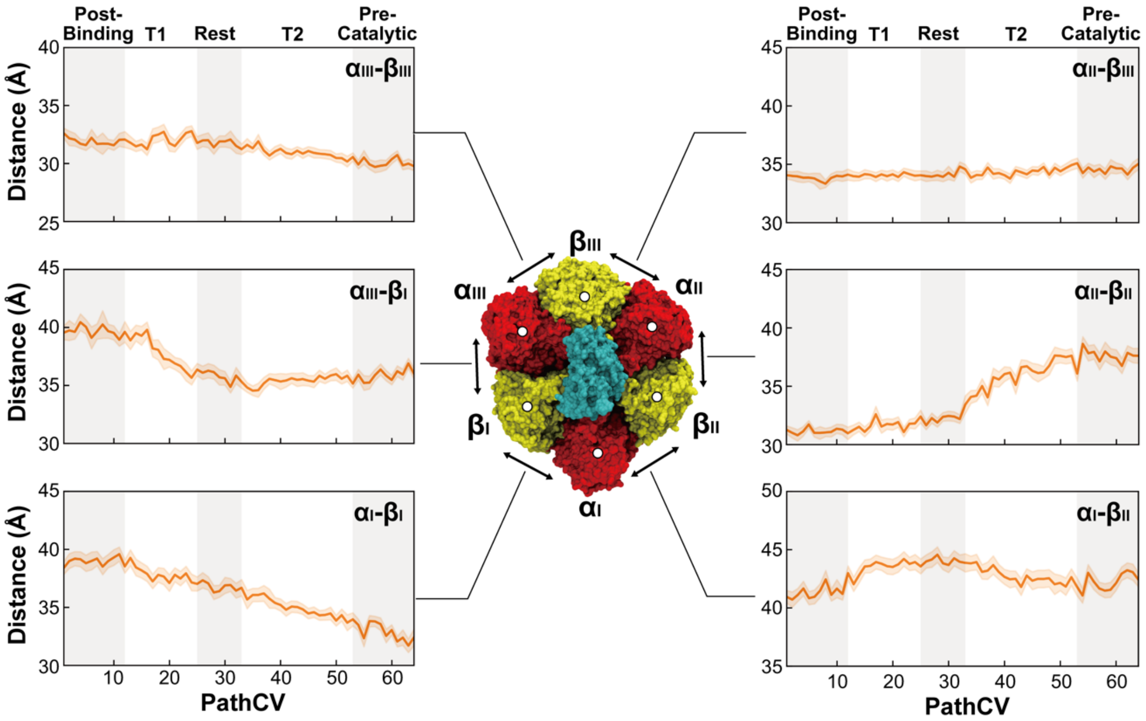


**Fig. S6. Distances between adjacent αβ interfaces during 80° rotation.**

The distances between the αβ interface are calculated from the centers of mass of Cα atoms in the C-terminal domain (CTD) of the α subunit (residues 371-499) and the CTD of the β subunit (residues 360-473). Data from 900 frames per window obtained from the US are used. The solid orange line in the graph represents the mean value, while the light orange shaded area indicates the standard deviation. White circles in the structural model represent the positions of the centers of mass.

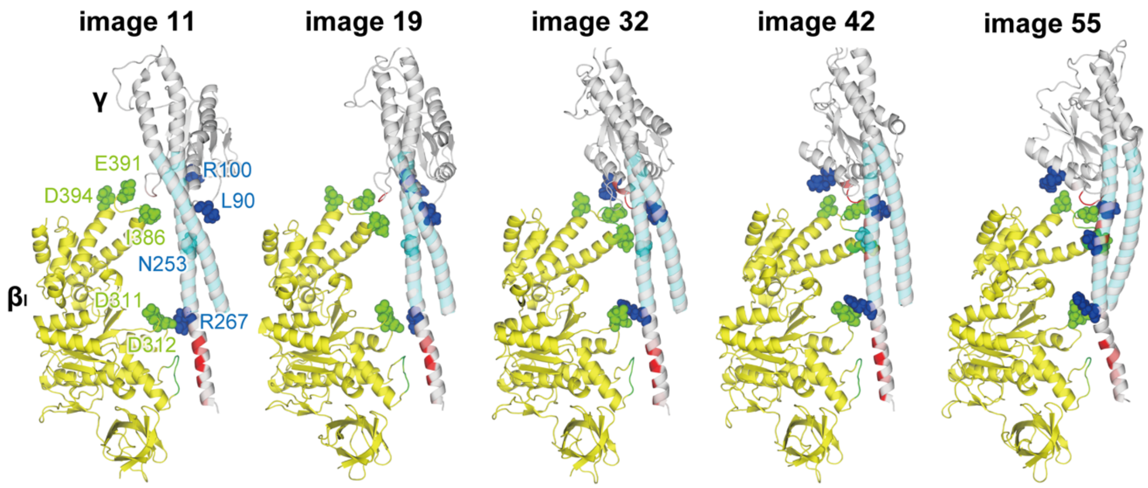


**Fig. S7. Interaction changes between $\beta_I$ and γ subunit.**

Representative structures of $\beta_I$ and γ in five distinct phases (images 11, 19, 32, 42, and 55) are shown. Residues in $\beta_I$ and γ that exhibit interaction changes are highlighted as green and blue spheres, respectively. Residues in γ and $\beta_I$ shown in cartoon representation and colored green and red indicate Cα atom distances between $\beta_I$ and γ of 8 Å or less.

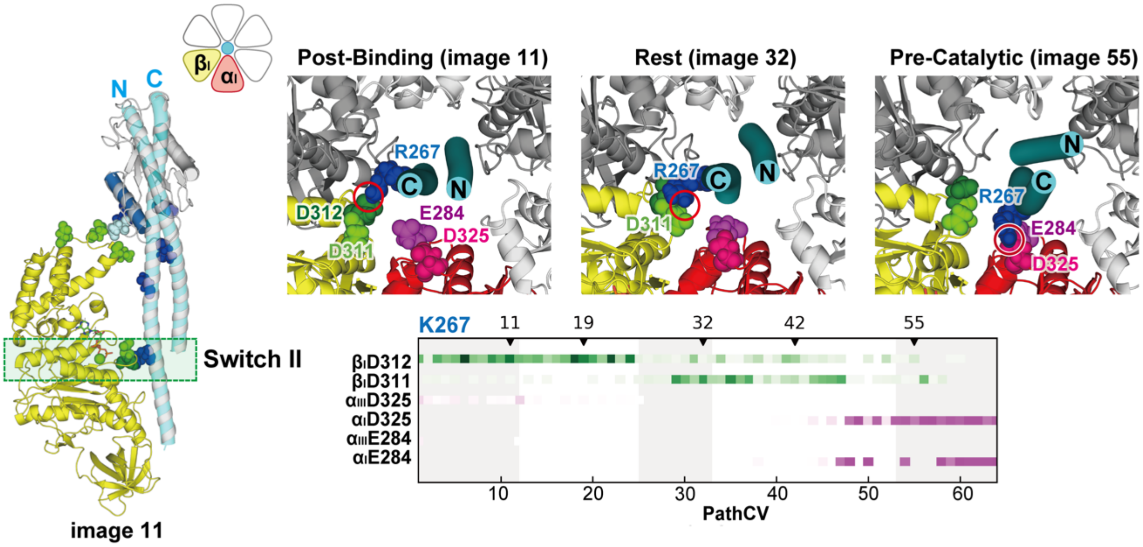


**Fig. S8. Interaction changes between $\alpha_I\beta_I$ and γ subunits in the switch II region.**

Residues within the α, β, and γ subunits that form the main interactions are highlighted as magenta, green, and blue spheres in images 11, 32, and 55, respectively. The following graph shows the frequency of interactions for each residue pair, with the high frequencies observed during the 90-ns umbrella sampling shown in dark colors.

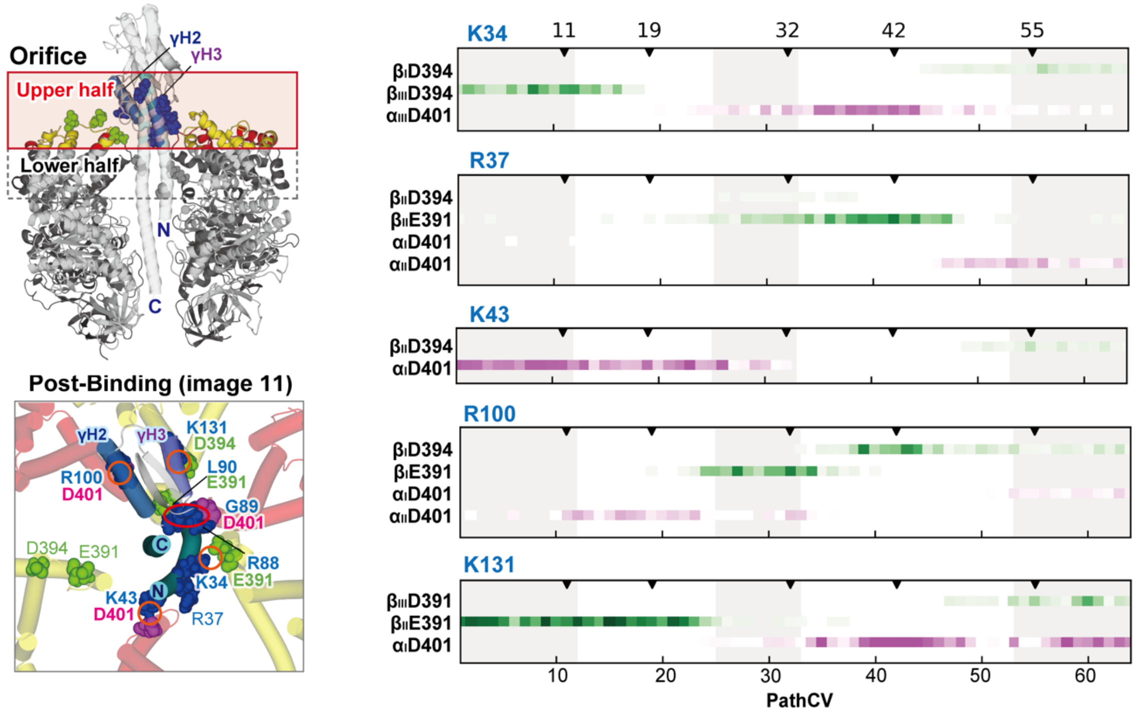


**Fig. S9. Interaction changes between $\alpha_3\beta_3$ ring and γ subunits in the upper half of the orifice region.**

Residues in the α, β, and γ subunits that form significant interactions between the upper CTD region of $\alpha_3\beta_3$ and the γ subunit are highlighted as magenta, green, and blue spheres, respectively, in image 11. The graph shows the change in the interaction between residues K34, R37, K43, R100, and K131 of the γ subunit and $\alpha_3\beta_3$. The change in the interaction of other residues (R88, G89, and L90) that form an interaction at the upper half of the orifice region is shown in Fig. 4B.

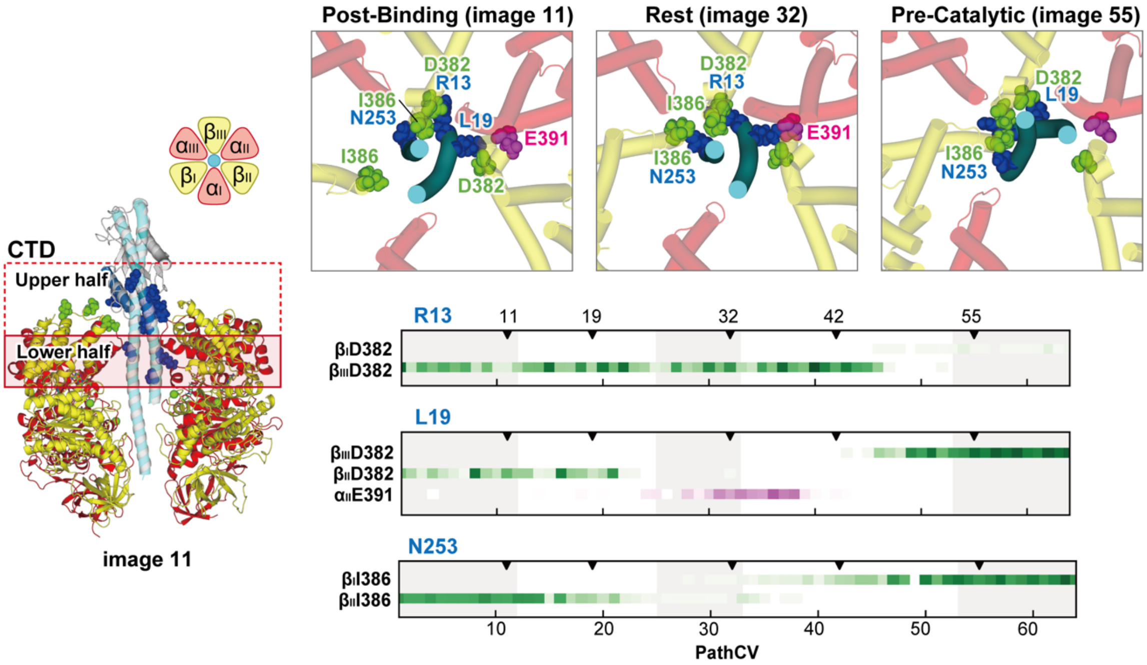


**Fig. S10. Interaction changes between α₃β₃ ring and γ subunits in the lower half of the orifice region.**

Residues in the α, β, and γ subunits that form significant interactions between the upper CTD region of $\alpha_3\beta_3$ and the γ subunit are highlighted as magenta, green, and blue spheres, respectively, in images 11, 32, and 55. The graph shows the change in the interaction between residues R13, L19, and N253 of the γ subunit and $\alpha_3\beta_3$.

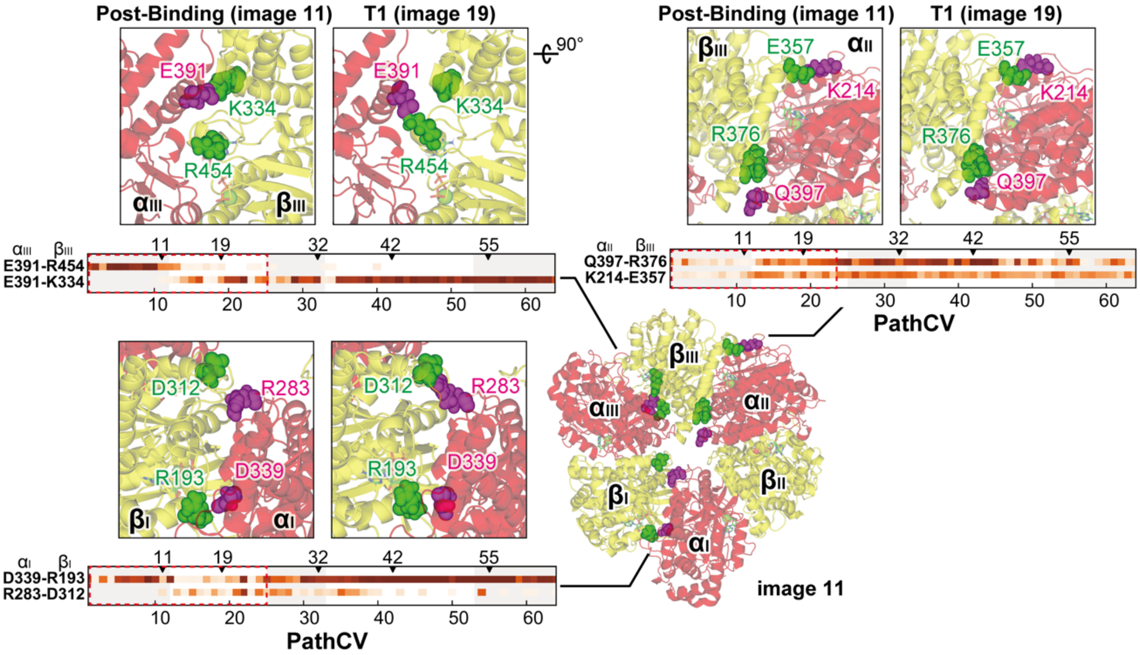


**Fig. S11. Interaction changes at αβ interfaces.**

Interactions formed and broken between α and β subunits during the transition are shown, and the changes from Post-Binding to T1 are highlighted with a red dotes square. In the graph, darker dots indicate more frequent interactions throughout the 90 ns US simulation.

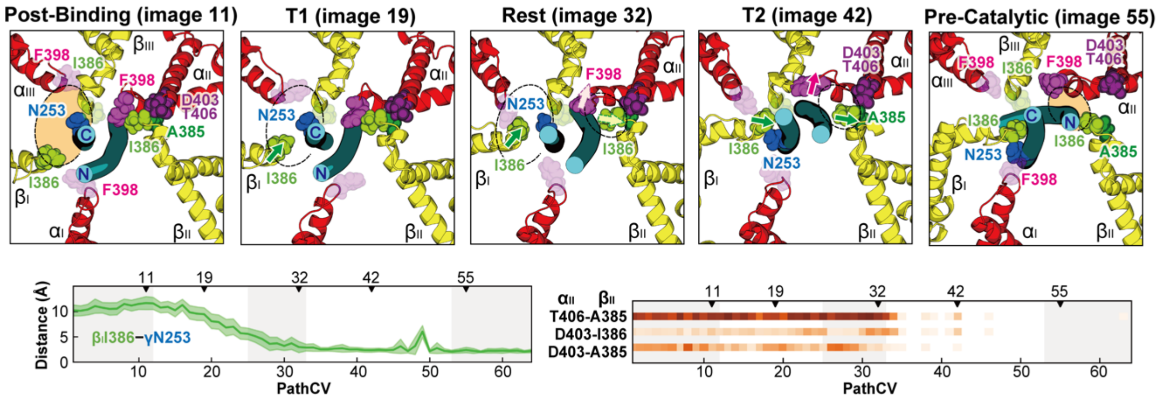


**Fig. S12. Contact between $\beta_I$ and γ and changes in the interaction at the $\alpha_{II}\beta_{II}$ interface in the lower part of the orifice region.**

Top view of the residues at the lower half of the orifice region is shown for images 11, 19, 32, 42, and 55. βI386 (light green) and αF398 in the CTD of the α and β subunits are highlighted as spheres. The graph on the lower left shows the interatomic distance between $\beta_I$I386 and γN253. The graph on the lower right shows the change in electrostatic interactions between $\alpha_{II}$ and $\beta_{II}$.

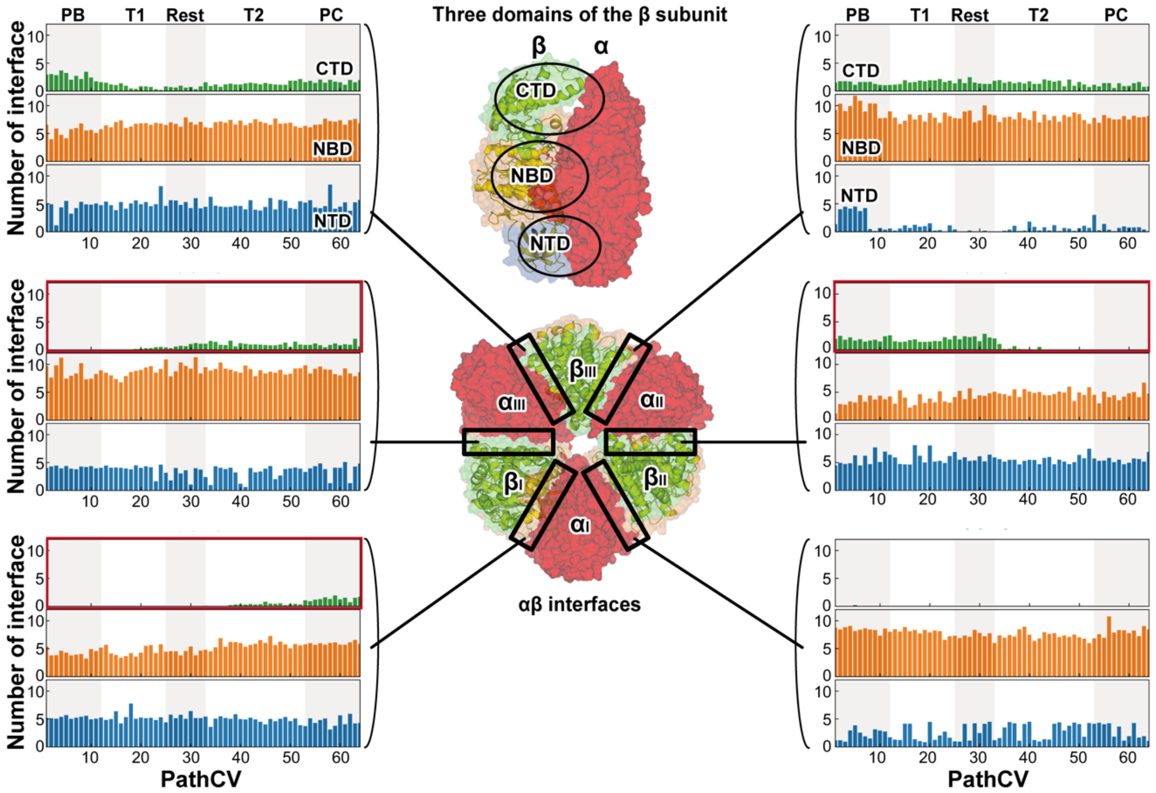


**Fig. S13. The number of electrostatic interaction changes between the three domains of the β subunit and the α subunit.**

Changes in the number of electrostatic interactions at the αβ interface. The interactions are counted separately for the N-terminal domain (NTD: residues 2-82, blue), nucleotide-binding domain (NBD: residues 83-359, orange), and C-terminal domain (CTD: residues 360-473, green) of the β subunit.

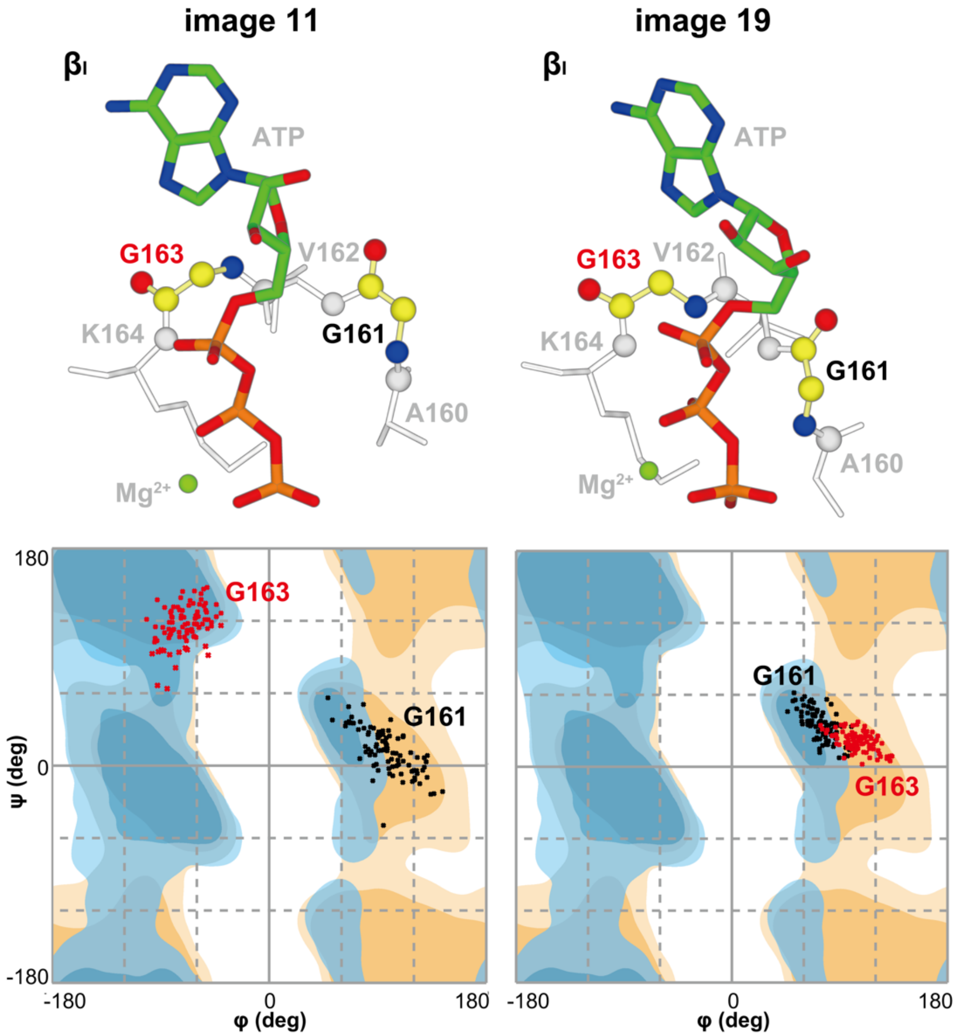


**Fig. S14. Dihedral angle changes of glycine in the P-loop.**

The first row shows the 3D structures of G161 (black) and G163 (red) within the $\beta_I$ P-loop in images 11 and 19. The second row displays Ramachandran plots for G161 and G163. Red and blue dots represent dihedral angle data points for G161 and G163, respectively, taken from 90 frames over the 90 ns US simulation. The reference dihedral angle map uses RAMPAGE's data (1), where blue and orange regions indicate the preferred (darker) and allowed (lighter) dihedral regions for general residues and glycine, respectively.

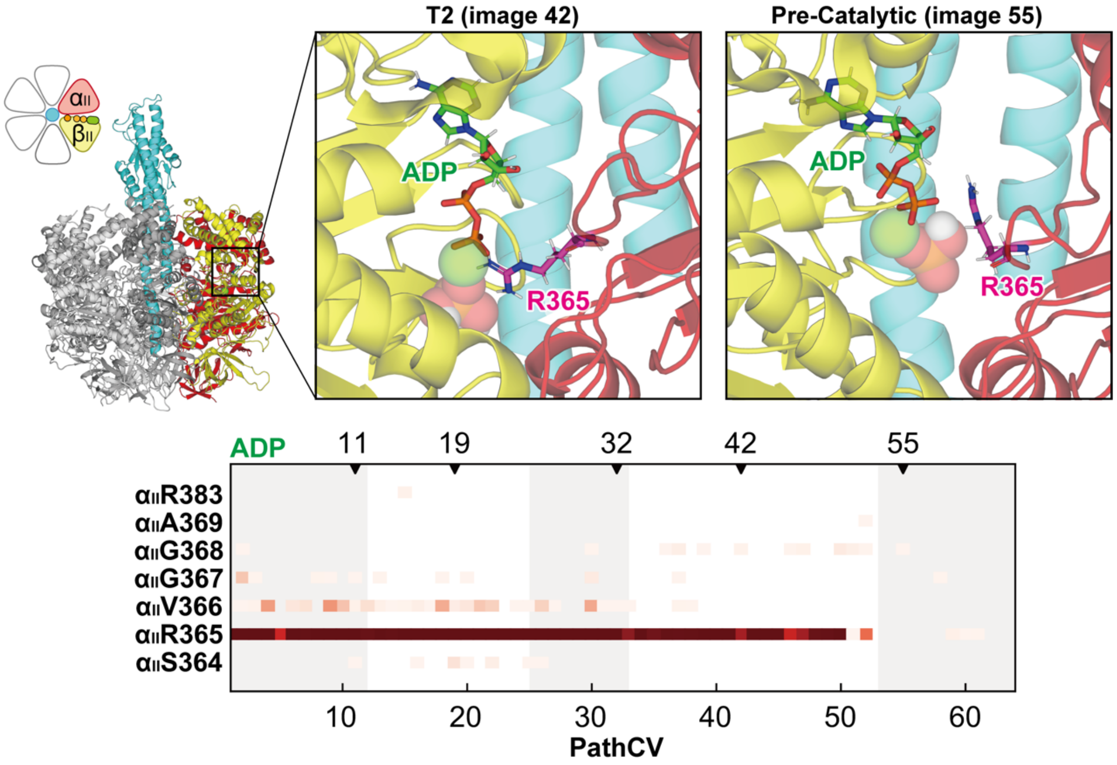


**Fig. S15. Electrostatic interaction changes between $\alpha_{III}$ and ADP.**

The changes in electrostatic interactions between $\beta_{III}$ and ADP are represented by shades of orange. The y-axis labels indicate the residue names and residue numbers of $\beta_{III}$. Dots with darker shades indicate higher interaction frequency throughout the 90-ns US simulation.

**Table S1.** Free energy changes with and without nucleotides. The table on the left shows the difference in free energy between Post-Binding (image 11) and T1 (image 19) depending on the presence or absence of $\beta_I$-bound ATP. The table on the right illustrates the difference in free energy between Pre-Catalytic (image 55) and T2 (image 42) based on the presence or absence of $\beta_{II}$-bound ADP.

| | Post-Binding – T1 $\Delta G_{11,19}$ (kcal/mol) |
|---|---|
| + $\beta_I$ATP | 9.75 |
| – $\beta_I$ATP | – 8.45 |

| | Pre-Catalytic – T2 $\Delta G_{55,42}$ (kcal/mol) |
|---|---|
| + $\beta_{II}$ADP | 4.55 |
| – $\beta_{II}$ADP | – 1.60 |

**Movie S1 (separate file).** The transition pathway of an 80° rotation obtained using the mean-force string method with 64 images. The CTDs of the α and β subunits are highlighted in red or yellow, while the other domains are shown in gray. The cyan dots in the upper left corner of the video indicate changes in the rotational angle of the γ subunit.
